# Comparing genome-based estimates of relatedness for use in pedigree-based conservation management

**DOI:** 10.1101/2021.07.08.451704

**Authors:** Samantha Hauser, Stephanie J. Galla, Andrea S. Putnam, Tammy E. Steeves, Emily K. Latch

## Abstract

Researchers have long debated which estimator of relatedness best captures the degree of relationship between two individuals. In the genomics era, this debate continues, with relatedness estimates being sensitive to the methods used to generate markers, marker quality, and levels of diversity in sampled individuals. Here, we compare six commonly used genome-based relatedness estimators (kinship genetic distance (KGD), Wang Maximum Likelihood (TrioML), Queller and Goodnight (R_xy_), Kinship INference for Genome-wide association studies (KING-robust), and Pairwise Relatedness (R_AB_), allele-sharing co-ancestry (AS)) across five species bred in captivity–including three birds and two mammals–with varying degrees of reliable pedigree data, using reduced-representation and whole genome resequencing data. Genome-based relatedness estimates varied widely across estimators, sequencing methods, and species, yet the most consistent results for known first order relationships were found using R_xy_, R_AB_, and AS. However, AS was found to be less consistently correlated with known pedigree relatedness than either R_xy_ or R_AB_. Our combined results indicate there is not a single genome-based estimator that is ideal across different species and data types. To determine the most appropriate genome-based relatedness estimator for each new dataset, we recommend assessing the relative: (1) correlation of candidate estimators with known relationships in the pedigree and (2) precision of candidate estimators with known first-order relationships. These recommendations are broadly applicable to conservation breeding programs, particularly where genome-based estimates of relatedness can complement and complete poorly pedigreed populations. Given a growing interest in the application of wild pedigrees, our results and are also applicable to *in-situ* wildlife management.

## Introduction

Relatedness and kinship, concepts that quantify the relationship between two individuals (hereafter referred to as relatedness for simplicity) are foundational in biology (Wright, 1922), with application in health sciences (Ott, 1974), agriculture (Cassell et al., 2003; Jannink et al., 2001) and species conservation (Fernández et al., 2005). Pedigrees track relatedness in populations by documenting the ancestry of individuals, and are used to manage small populations, including those in captivity (i.e., *ex situ* population management) or intensively managed wild or semi-wild populations (i.e., *in situ* or “*sorta situ”;* Wildt et al., 2019; Wolfe et al., 2012; Galla et al. 2022). Conservation practitioners can prioritize breeding individuals with low mean kinship when making pairing decisions in captivity, or population management decisions in wild or semi-wild populations (Ballou & Lacy, 1995; Giglio et al., 2016; Weeks et al., 2011). A pedigree-based management strategy that minimizes mean kinship is effective at mitigating drift, inbreeding, and adaptation to captivity, while preserving genetic diversity and evolutionary potential to curtail extinction risk (Fernandez & Toro, 1999; Montgomery et al., 1997; Sonesson & Meuwissen, 2001; Spielman, 2004, Templeton & Read, 1994). Simulation studies have shown that pedigree-based management of small populations minimizes inbreeding while maximizing founder representation (e.g., Ballou & Lacy, 1995; Rudnick & Lacy, 2008). Empirical studies have demonstrated the importance of pedigree-based management for threatened species conservation (e.g., Tasmanian devil, *Sarcophilus harrisii*, Gooley et al, 2017; Ālala, *Corvus hawaiiensis*, Flanagan et al., 2021; Bison, *Bison bison*, Giglio et al., 2018). Because the collection of pedigree data is feasible for many captive and intensively managed wild populations, and computer programs are readily available for pedigree-based management (e.g., PopLink, ZIMS; Faust et al., 2019, Species360), pedigree analysis (e.g., PMx; Lacy et al., 2012), and population modeling (e.g., VORTEX; Lacy & Pollak, 2014), pedigree-based management of populations is an accessible and appealing tool for conservation programs.

Pedigrees continue to provide high precision estimates of inbreeding and relatedness when pedigrees are robust (i.e., many generations deep with relatively low missing data; Putnam & Ivy, 2014; Robinson et al., 2013). However, there are several limitations common in pedigrees for small populations that can hinder their utility for genetic management (Galla et al. 2022). One is that the individuals forming the initial captive population (hereafter, founders) are often of unknown relationships and assumed to be equally unrelated (Ballou 1983; Hogg et al., 2019). In small populations that have experienced sustained bottlenecks, it is unlikely that remaining individuals are unrelated. For example, a genetic study of critically endangered kākāpō*(Strigops habroptilus)* revealed high relatedness coefficients—including some first-order (i.e., parent-offspring or full sibling) relationships—among founders that were previously assumed to be unrelated (Bergner et al., 2014). This can be magnified when additional founders are incorporated into pedigrees. For example, when captive populations are augmented using wild individuals with unknown relationships to each other and to the captive individuals already in the population (Galla et al., 2020; Spielman & Frankham, 1992). Assuming founders are unrelated can lead to underestimated kinship and inbreeding coefficients, which is exacerbated when pedigrees are shallow (< 5 generations deep; Balloux et al., 2004 but see Rudnick and Lacy 2008).

Pedigree gaps indicating unknown relationships can also be a source of ambiguity constraining the utility of pedigrees for genetic management. Missing data can arise from uncertainties in parentage, for example through undetected extra-pair parentage or herd or colonial breeding systems (Overbeek et al., 2020; Ferrie, 2017; Mucha & Windig, 2009). In the wild, missing data may be introduced, or relationships incorrectly inferred due to individual identification errors (e.g., dropped leg bands; Milligan et al., 2003). Parentage assignment errors may be rare (Henkel et al. 2012, Ferrie et al. 2013, Hammerly et al. 2016), but their effects can compound over generations, thereby increasing the probability that related individuals will be unintentionally paired. For example, in a captive population of Attwater’s Prairie-chicken (*Tympanuchus cupido attwateri*), nearly 40% of the population were descendants of four individuals that had incorrect parentage assigned four generations prior (Hammerly et al., 2016). To alleviate pedigree shortcomings, including unknown founder relationships, missing pedigree data, and potential error, researchers can use genomic estimates of relatedness to complement and complete pedigrees (Galla et al., 2022).

Relatedness estimates generated using genomic markers have helped to resolve founder relationships (Bergner et al., 2014), reconstruct parentage (Flanagan & Jones, 2019), and validate pedigrees for potential errors (Hammerly et al., 2016). Factors that can affect the accuracy of genome-based relatedness estimates include marker type, number of loci, missingness, sequencing depth, and the specific estimator used. Single nucleotide polymorphisms (SNPs) - derived from SNP-assays (Santure et al., 2010), reduced representation sequencing (RRS) approaches like RAD-sequencing or GBS (Galla et al., 2019; Lemopoulos et al., 2019), or whole genome resequencing (WGS; Galla et al., 2020) yield accurate genomic estimates of relatedness (Jones & Wang, 2010; Santure et al., 2010b; Skare et al., 2009). However, the number of markers necessary for a given system depends upon the genome size of the species, population size, and overall level of inbreeding in the population; insufficient marker power can lead to inaccurate relationship inference (Goudet et al., 2018; Morin et al., 2009; Smouse, 2010; Sun et al., 2016). Similarly, the specific estimator used to capture inheritance patterns and translate them into a relatedness estimate can impact the accuracy of relationship inference.

There are several commonly used relatedness estimators appropriate for genomic data, each with its own limitations (Box 1). Briefly, frequency-based estimates of relatedness quantify the probability that shared alleles are identical-by-descent (IBD) relative to a reference population using probabilities (i.e., moment-methods) or correlations (likelihood-based; Wang, 2014). Frequency-based estimators assume that populations are large, randomly mating, and outbred (but see Hedrick & Lacy, 2015; Wang, 2007), effectively sampled (Wang 2017), and that allele frequencies at each marker are reliably estimated (Csillery et al. 2006, Galla et al. 2020). However, there are notable exceptions (see Box 1). A meaningful reference population for frequency-based relatedness estimation may be difficult to define, especially for small conservation breeding populations with non-random mating. Further, the precision of frequency-based approaches will rely on marker completeness and richness, with more markers leading to more precise estimates of relatedness (Baruch & Weller, 2008; Morin et al., 2009). As an alternative, pairwise allele sharing (i.e., molecular co-ancestry or similarity index; Gutiérrez et al., 2005) does not rely upon reference population allele frequencies. Further, Ivy et al. (2016) found allele-sharing to be strongly correlated with mean kinships derived from pedigrees using genomic data, thus making it a potentially useful estimator for populations where the assumptions of frequency-based estimators are violated, such as those in conservation breeding programs.

Best practice guidance to estimate accurate genome-based relatedness using high throughput sequencing data is needed to maximize conservation success for poorly pedigreed threatened populations. Examining the relative performance of genome-based estimators using empirical data can expose estimator pitfalls that cannot be found solely through simulation studies. In this study, we extend previous empirical studies of relatedness estimators (Attard et al., 2017; Csillery et al., 2006; Goudet et al., 2018, Santure et al., 2010) to address persistent challenges to genome-based relatedness estimation for intensively managed non-model species. Briefly, we compare the accuracy of six genome-based relatedness estimators relative to observed pedigree data, including allele-sharing and five different frequency-based estimators across 9 genomic datasets. Relatedness estimates were derived from SNP data generated using at least one of three different approaches (WGS, ddRAD or GBS), collected from five different species with *ex-situ* programs, including Addax (*Addax nasomaculatus*), Inca Tern (*Larosterna inca*), Koala (*Phascolarctos cinereus*), Kakī (Black Stilt; *Himantopus novaezelandiae*) and Kākāriki Karaka (Orange-fronted Parakeet; *Cyanoramphus malherbi;* Table 1). In addition to several interactive factors related to the data itself (i.e., genome size, inbreeding, marker type, number of loci, missingness, sequencing depth), these five species offer a variety of realistic challenges in conservation breeding programs, including the size of the managed population, depth of the pedigree, and the amount and structure of pedigree gaps (Figure 1). Beyond broad applicability to conservation breeding programs, pedigree-informed management of wild populations (Pemberton, 2008) will benefit from explicit evaluation of genome-based relatedness estimators.

**Figure 1.**
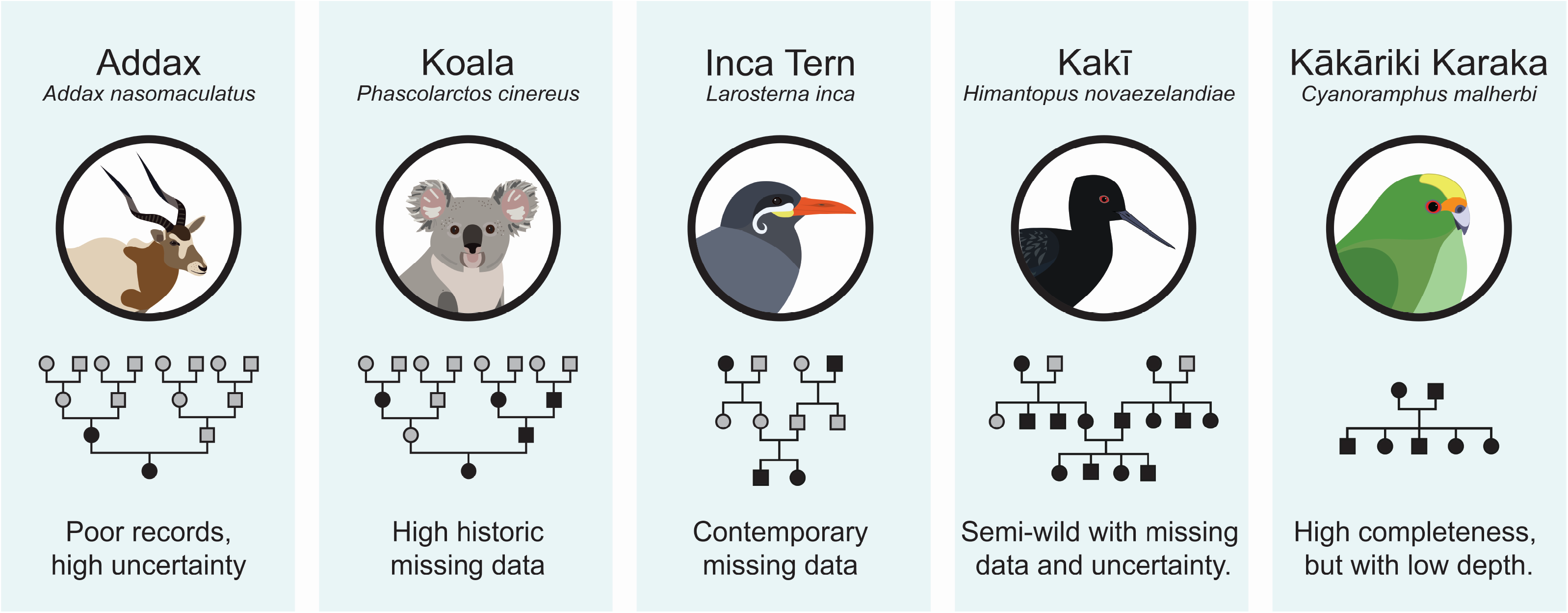
Key pedigree challenges faced by the conservation breeding programs for each of the species used in this study. Schematic pedigrees denote missing data with white circles/squares and existing data with black circles/squares.

**Table 1.**
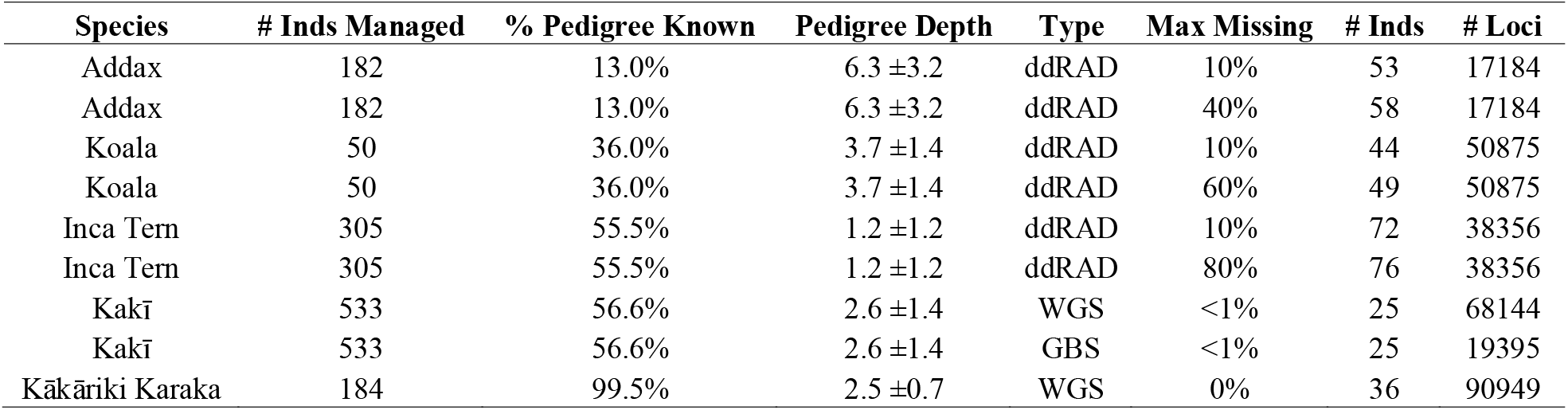
Summary information on each of the 9 genomic datasets used including: Number of individuals currently managed in the captive population (# Inds Managed), percentage of the pedigree known prior to genomic data incorporation (% Pedigree Known), average generation per individual with standard deviation (Pedigree Depth), type of genomic technique used (Type), the maximum missingness threshold used (Max Missing), number of individuals with SNP data in the dataset (# Inds), number of SNP loci in the dataset ( #Loci), percentage of the pedigree known after genomic data incorporation (% analytic known)

## Methods

Of the five species with breeding programs in the present study, three (Addax, Inca Tern and Koala) populations are cooperatively managed through the Association of Zoos and Aquariums (AZA) Species Survival Plans^®^, and two (Kakī and Kārāriki Karaka) are managed by, or on the behalf of, the New Zealand Department of Conservation.

### Addax

The current population of addax in AZA descends from a small number of individuals (~15) imported from Chad and the Khartoum Zoo, Sudan in the 1960s and 1970s (Enright, 2019). Until the early 2000s, facilities with large herds did not reliably record parentage and frequent exchanges with the private sector occurred. At the time of analysis, only 13% of the AZA addax population could be traced to specific founders.

### Koala

The contemporary pedigree for the AZA Queensland koala population is well recorded. However, unresolved relationships in the historical pedigree of captive-bred Australian importations, mostly from a single facility in the 1970s – 1980s, generates an overall pedigree for the living AZA population that is only 39% is known (Hamlin, 2019).

### Inca Tern

Wild Inca terns were imported by AZA facilities in large numbers during the 1960s through the mid-1990s. Parentage has not been reliably recorded at one facility that holds and propagates the largest number of terns within the AZA, resulting in a steady increase in the proportion of the pedigree that cannot be traced to founders (30%; Nelson, 2017, Nelson and Lynch, 2021).

### Kakī

Kakī are a critically endangered wading bird endemic to the South Island of Aotearoa New Zealand. While Kakī were once more widespread in Aotearoa, they experienced decline due to introduced mammalian predators and habitat loss (Sanders & Maloney, 2002). By the early 1980s, only 23 birds remained (Galla, 2020). Since then, intensive management strategies—including predator control efforts and a conservation breeding program—were initiated to prevent extinction and enhance recovery. The Kakī pedigree has been maintained for this population and includes intensively managed wild and captive individuals. About half of the Kakī pedigree is known (56.6%). Pedigree gaps come from unbanded birds (gaps) and unaccounted extra-pair copulation (errors) in the wild (Overbeek et al., 2020).

### Kākāriki Karaka

Kākāriki Karaka were once more widespread on the South Island of Aotearoa, and experienced decline due to invasive mammalian predators and habitat loss. During the 20^th^ century, this species was considered extinct twice, likely due to its cryptic nature. In 2003, a conservation breeding program was initiated with 12 individuals collected from the wild, with sparse supplementation in recent years (Galla, 2020). The Kākāriki Karaka pedigree is a nearly complete pedigree, with 99.5% of the pedigree known. While complete, this pedigree is shallow (0-3 generations deep), and as a result may be more impacted by incorrect assumptions about founder relationships.

We generated SNPs for each of the five species using two types of sequence data, RRS or WGS (Table 1): double-digest RAD (ddRAD) sequencing data for the Addax, Inca Tern, and Koala (herein), GBS data for Kakī (Galla et al., 2019), and WGS for Kakī and Kākāriki Karaka (Galla et al., 2020). The bioinformatic methods and parameters selected for SNP discovery and genotyping vary as they were chosen based on best practice optimization protocols for each sequencing method and the type of empirical data observed for each species.

We isolated high quality DNA from blood, feather, or tissue samples for Kakī and Kākāriki Karaka using a lithium chloride chloroform extraction method (Galla et al., 2019) and quantified using a Qubit 2.0 Fluorometer. Kakī GBS paired-end libraries were prepared using the Illumina TruSeq DNA PCR-free protocol (350 bp DNA fragment) and sequenced on 5 lanes of an Illumina HiSeq 2500 (Galla et al., 2019). Kakī WGS libraries were prepared using the TruSeq Nano DNA Library Prep Kit and sequenced on 34 lanes of Illumina HiSeq 400 (Galla et al., 2020) Kākāriki Karaka WGS libraries were prepared using the Nextera DNA Flex Library Prep Kit and sequenced on 1 lane of an Illumina Novaseq 6000 (Galla et al., 2020). SNP data from WGS and GBS data for Kakī and Kākāriki Karaka were processed, and genotypes were called as per Galla et al., (2019, 2020). Briefly, Kakī GBS data was generated from two batches of individuals, Illumina Hi-Seq reads underwent reference-guided SNP discovery using Tassel 5.0 (Glaubitz et al., 2014), and biallelic SNPs were filtered for a minimum MAF of 0.05, an average minimum SNP-depth per locus of five, maximum of 10% missing data, and linkage disequilibrium (*r^2^* = 0.6 over 1000 sites; Galla et al., 2019). For Kakī and Kākāriki Karaka WGS, reads were generated from Illumina Hi-Seq or NovaSeq platforms, SNPs were discovered using BCFtools v. 1.9 (Li et al., 2009), and biallelic SNPs were filtered for a minimum MAF of 0.05, a quality score greater than 20, a maximum of 10% missing data, linkage disequilibrium (*r^2^* = 0.6 over 1000 sites), and either a minimum depth of 5 (Kākāriki Karaka) or a minimum average depth of 10 (Kakī; for more details, see Galla et al., 2020). GBS and WGS Kakī data were from the same consensus 25 individuals to allow direct comparisons between SNP data generation approaches (Table 1).

We extracted high quality DNA from blood samples for Inca Tern, Koala, and Addax using the Qiagen DNeasy Blood and Tissue Kit. All DNA isolates were quantified using a Qubit 2.0 Fluorometer. For the three AZA species, ddRAD library preparation (per Peterson et al., 2012), quality control, and sequencing were done at the Texas A&M AgriLife Genomics core facility. ddRAD libraries were prepared using restriction enzymes *Spe*I and *Mbo*I (Koala and Addax) or *Sph*I and *MluC*I (Inca Tern) for paired-end 150 bp reads and sequenced on a portion of an Illumina NovaSeq 6000 lane. Raw sequencing data were demultiplexed, filtered, and genotypes were called using the bioinformatics pipelines STACKS v.2.0 and VCFTOOLS (Catchen et al., 2013; Danecek et al., 2011). SNP genotypes for Addax and Inca Tern were generated using the de novo pipeline in STACKS with the following parameters chosen through optimization (Paris et al., 2017), respectively: m = 3, M = 3, n = 0, min_maf = 0.02, r = 0.7 and m = 3, M = 3, n = 0, min_maf = 0.02, r = 0.6. Genotypes were called for Koala using the reference pipeline using a reference genome (GenBank Accession: GCA_002099425.1) with min_maf = 0.02 and r = 0.80 parameters. Individuals were omitted using VCFTOOLS based on maximum missingness thresholds, set to two settings for each dataset to quantify missing data impacts: low (10% for all datasets) and high (staggered at 40% in Addax, 60% in Koala, and 80% in Inca Tern; Table 1). In conservation breeding programs, a combination of unavoidable suboptimal samples due to sampling regulations (e.g., mammalian blood) and low species-wide genetic diversity can lead to high missing data (Graham et al., 2015). Low genetic diversity does not beget high missing data, but exacerbates high missing data associated with suboptimal sample quality. As such, we tested several threshold filters for missing data to encapsulate the variety of high missing datasets likely to burden conservation breeding programs. Maximum values for the high missing datasets were chosen based on the observed distribution of missing genomic data found for each species.

These processing steps yielded nine SNP datasets–Addax, Koala and Inca Tern each with a low missing data and a high missing data ddRAD dataset, Kakī with a WGS and GBS dataset and Kākāriki Karaka with a WGS dataset – for which we calculated six genome-based relatedness estimators: Allele-Sharing (AS), Kinship Genetic Distance (KGD; Dodds et al., 2015), Wang Maximum Likelihood (TrioML; Wang, 2007, 2011), Queller and Goodnight (R_xy_; Queller & Goodnight, 1989), Kinship INference for Genome-wide association studies (KING-robust; Waples et al., 2019), and R_AB_ (Korneliussen & Moltke, 2015; Box 1). The AS estimates were calculated in the program CASC (Ivy & Putnam, 2019), subsampling 1,000 loci and using 1,000 iterations. AS values are scaled differently than genome-based kinships, so we transformed AS values using a linear piecewise regression on known pedigree-based kinship estimates (>0).

Pedigree-based kinship estimates were calculated in PMx (Lacy 2012) based on PopLink studbook data and were scaled to relatedness estimates (2*kinship) for comparative analyses. KGD estimates were calculated in R (R Core Team, 2018) using the approach described in Dodds et al. (2015). TrioML and R_xy_ estimates were calculated using the R package ‘related’ (Pew et al. 2015), specifying the settings ‘unknown allele frequencies’ (i.e., allele frequencies are calculated from the imported data) and ‘inbreeding’. KING-robust and R_AB_ estimators were estimated using NGSRelate (Hanghøj et al, 2019). All kinship (KING-robust) estimates (scale = 0 – 0.5) were multiplied by 2 to approximately scale them to the rest of the genome-based relatedness estimators and pedigree relatedness values (scale = 0 – 1.0). As transformations were necessary to directly compare genome-based relatedness estimators, we checked all estimator distributions pre- and post-transformation to ensure that our transformations were not distorting the data.

We compared estimator performance across all nine SNP datasets in two ways. First, each pairwise relatedness estimator was directly compared to known pedigree relatedness and compared to all other genome-based relatedness estimators using Pearson’s correlation coefficients in R. For these analyses, only those individuals with some degree of known pedigree relatedness (i.e., those with relatedness values > 0) were included. Second, parent-offspring, full sibling, and half sibling relationships were derived from each dataset, and boxplots of each relatedness estimator and pedigree relatedness value were plotted in R package *ggplot2* (Wickham, 2011). The only exceptions were the two Kakī datasets which lacked half siblings. We included both first-order and second-order relationships to contrast our expectations: whereas both genome-based relatedness estimators and pedigree-based relatedness values for first-order relationships (parent-offspring, full sibling) are anticipated to be relatively robust with low expected variance–making them ideal to explicitly test the relative performance of genome-based relatedness estimators–variance in both genome-based relatedness estimators and pedigree-based relatedness values is expected to be larger for more distant relationships (e.g., half siblings).

## Results

We generated six sets of genome-based relatedness estimates for each of the nine genomic datasets (Table 1). Rxy (mean Pearson’s *r* = 0.82, variance = 0.004), followed by R_AB_ and TrioML (mean Pearson’s *r* = 0.81, variance = 0.003) were the most consistently and highly correlated with pedigree data across all species and datasets (Figure 2). The least consistent estimators across all datasets were KGD (variance in Pearson’s *r* = 0.036) and KING-robust (variance in Pearson’s *r* = 0.022). Lower correlation values in certain estimators (i.e., KGD and KING-robust) were found in both low and high missing genomic datasets (Addax 10 and Addax 40, Inca Tern 10 and Inca Tern 80; Figure 2), which could suggest that other factors, not represented in our comparisons, may be influencing estimator performance. The most consistently high correlation values between genome-based and known pedigree-based relatedness values were found in datasets with low missing data and a large number of loci. Higher correlations were also found in WGS data (Pearson’s *r* = 0.69 – 0.88) than in GBS data (Pearson’s *r* = 0.4 – 0.79), which had considerably fewer loci and thus lower resolution (Kakī WGS = 68,144 and GBS = 19,395). In general, datasets with more SNPs produced estimates of relatedness with higher correlation with known pedigree relatedness than datasets with smaller numbers of SNPs (Figure 2). This pattern was particularly observed when comparing WGS and GBS datasets in Kakī (WGS mean Pearson’s *r* = 0.81; GBS mean Pearson’s *r* = 0.70; Figure 2).

**Figure 2.**
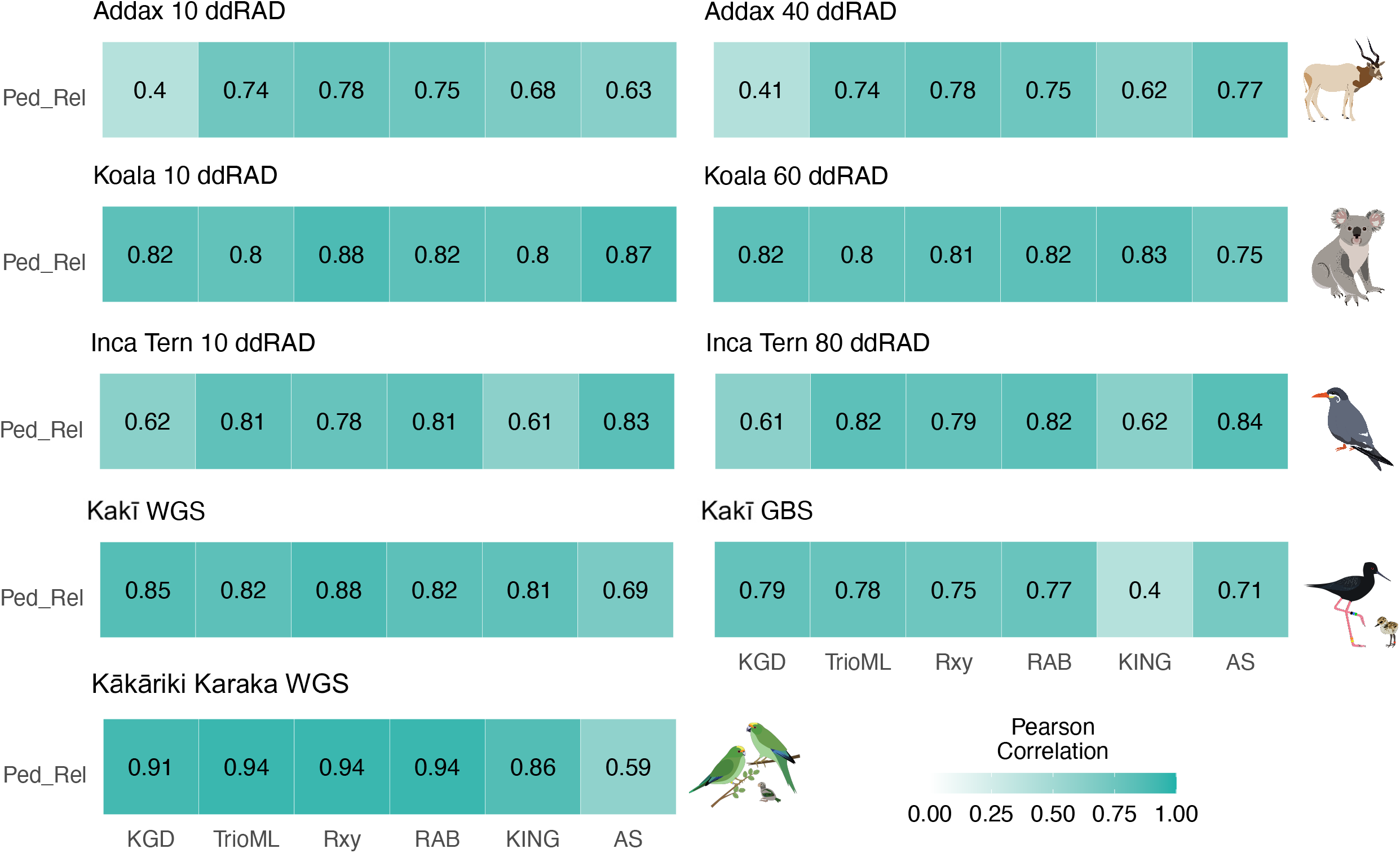
Heatmaps depicting pairwise Pearson’s correlation coefficients between pedigree relatedness (Ped_Rel) and six relatedness estimators for each of the genomic datasets used in this study with darker gradient colors denoting higher correlations. See text for details.

Genome-based relatedness estimators underestimated close pedigree relatedness values across species, missing data, and sequencing type (Figure 3). All pedigree estimates of first order relationships showed little variation, except for kakī, which has more intergenerational inbreeding not accounted for by our kinship duplication (*data not shown*). Across species, the precision of genome-based estimators varied widely. Estimators were comparable in the Inca Tern and Koala datasets but varied widely for Addax and Kākāriki Karaka. The relative performance of estimators was generally consistent between low and high missing data counterparts (i.e., Addax 10 and Addax 40, Koala 10 and Koala 60, Inca Tern 10 and Inca Tern 80; Figure 3). While there are some differences in the boxplot distributions between missing data counterparts such as slightly wider ranges (e.g., AS in Addax), different medians (KING-robust in Koala), or varied outliers (e.g., RAB in Inca Tern) in the high missing datasets. Between sequencing approaches, i.e., Kakī WGS and GBS, we found wider distributions and a large difference with the boxplot distributions of KING-robust (Figure 3).

**Figure 3.**
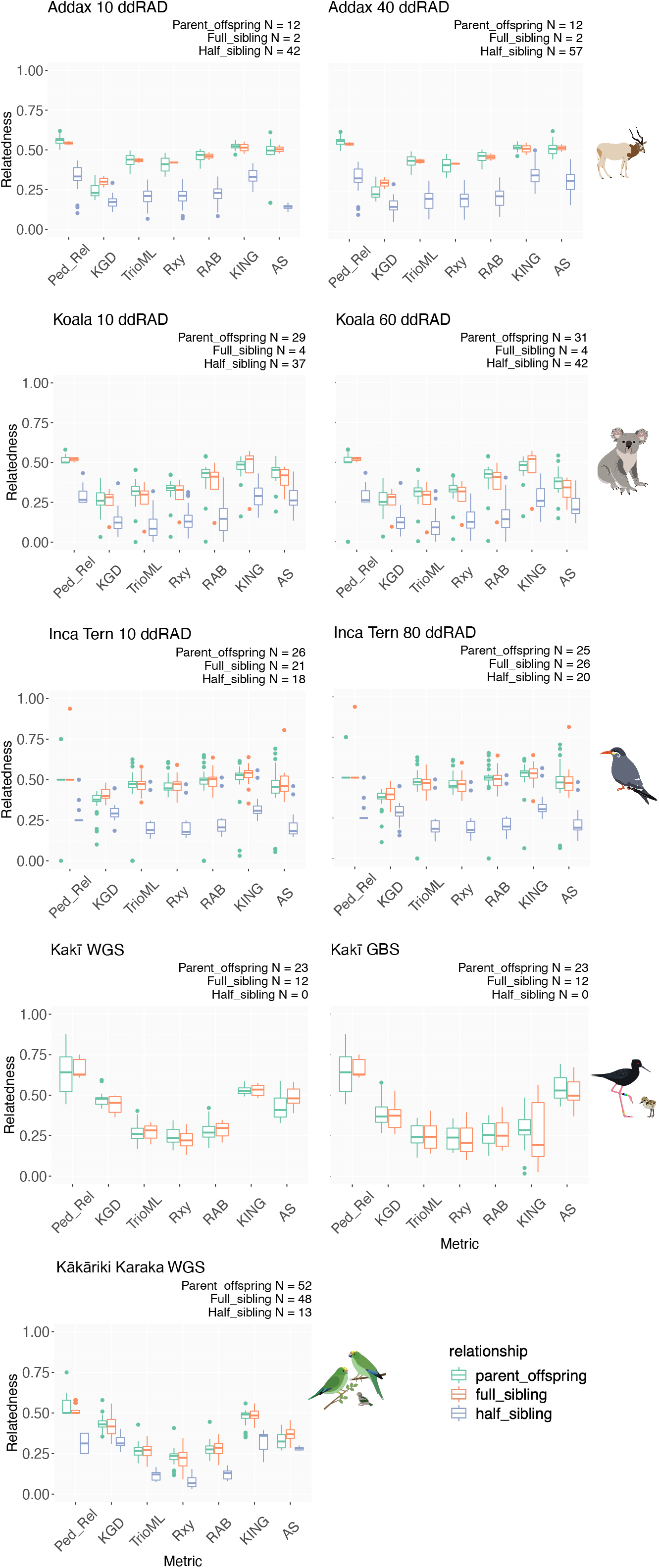
Boxplots of pedigree relatedness and six relatedness estimators of known relationship classes (parent-offspring (orange), full sibling (green), and half sibling (blue)) for each of the species used in our study. Relatedness values from 0 - 1 are on the y-axis and associated relatedness estimator names are on the x-axis.

## Discussion

Evaluation of relatedness estimators is a necessary step prior to the incorporation of genomic data into conservation breeding programs. Attributes of the genomic dataset and the underlying pedigree impact the performance of relatedness estimators in different ways, and the optimal estimator varies across datasets. Theoretical expectations assert that allele-sharing would be a suitable genome-based relatedness estimator for intensively managed conservation breeding populations, given that it does not require a reference population and is shown to be well correlated with pedigree relatedness estimates (Ivy et al., 2016). While allele-sharing estimators without a transformation are inappropriate for use, our results show that transformed allele-sharing values perform comparably to the other relatedness estimators. Because allele-sharing provides a direct measure of identity-by-state, as opposed to a frequency-based estimate of identity-by-descent, this estimator is expected to overestimate pairwise relatedness when genetic diversity or more specifically minor allele frequencies are low. This is salient for estimating relatedness in genetically-depauperate conservation breeding programs, where the probability of identity-by-state is inflated (Henkel et al., 2012). As a result, allele-sharing values are on a vastly different scale than pedigree relatedness and frequency-based relatedness estimators and any potential use of allele-sharing requires rescaling using a complex transformation for incorporation into pedigree-based management, for example, the linear piecewise transformation used here. While allele-sharing estimators without a transformation are inappropriate for use, our results show that transformed allele-sharing values perform comparably to the other relatedness estimators. However, it is important to note that because these transformations used existing pedigree data to rescale the allele-sharing data, the correlations may inherently bias correlations upward. Pedigree data may also not be available to transform raw allele-sharing data to the appropriate scale across systems. Further, transformations can introduce errors and reduce accuracy and precision of relatedness estimates; future use of allele-sharing should use caution in reducing potential errors and accounting for errors from the transformation in their interpretations.

We anticipated that the inability to accurately estimate population allele frequencies in conservation breeding programs would lead to biased relatedness estimates when using frequency-based estimators. Although all genome-based estimators consistently underestimated relatedness across all datasets, we generally found high correspondence between relatedness estimators and known pedigree values, especially in cases of low missing data, more fully resolved pedigrees, and high-resolution genomic data (WGS and large numbers of SNPs).

Confounding variables across empirical datasets (e.g., pedigree accuracy, degree of inbreeding, number of individuals, species genome size) affect relatedness estimates and contribute to inconsistent estimator performance across datasets. The advantages and caveats unique to each relatedness estimator means that there is no single optimal estimator for all datasets. Therefore, we recommend that researchers formally test candidate estimators for their systems by investigating the relative correlation with known relationships in the pedigree (e.g., Figure 2) and the relative precision of candidate estimators for known relationship categories (e.g., Figure 3). Estimators that do not approximate known parent-offspring or full sibling relationships may not be appropriate to for incorporation into identity-by-state pedigrees. Low correlations between known pedigree-based relatedness values and genome-based relatedness estimators likely indicate an unsuitable estimator for the data. If low correlations are found across a variety of estimators that use different approaches, the data itself may be insufficient for relatedness estimation, for example due to sampling deficiencies or sequence data that is low quality (missing data) and/or low quantity (few SNPs).

Missing data can occur in genomic datasets generated for conservation breeding populations for several reasons. While proactive collection and banking of high-quality DNA samples for genomic analyses is ideal to mitigate many of the limitations due to missing data, logistical and ethical constraints in handling captive animals (i.e., minimizing handling and the associated stress and risks) makes collection of high-quality and representative samples a challenge. These limitations often compel researchers to rely on low-quality samples opportunistically taken from veterinary samples including mammalian blood or plasma samples. Missing data can be the direct result of suboptimal quality samples as genetic material is low yield and/or degraded (Graham et al., 2015). A secondary effect of opportunistic sampling is that sequencing is often done in batches as samples become available through routine veterinary visits. Batch effects can be common and potentially problematic, reducing the number of loci shared among sequence batches and subsequently increasing the amount of missing data in a genomic dataset (Leigh et al., 2018). Thus, proactive sampling and storage of high-quality samples is important in reducing the downstream complications that can arise from opportunistic sampling (i.e., missing data). Conservation breeding managers can learn from ongoing efforts to establish proactive and standardized sample collection by groups including San Diego Zoo Wildlife Alliance’s Frozen Zoo (e.g., Chemnick et al., 2009), Cryo-Initiative (Comizzoli, 2014), Frozen Ark (Clarke, 2009), and the European Association of Zoos and Aquarium’s Biobank (see: https://www.eaza.net/conservation/research/eaza-biobank/).

Pedigreed populations will undoubtedly benefit with the addition of genome-based relatedness estimators. For pedigrees with little information regarding founders, or a large number of unknown relationships, the addition of genome-based relatedness estimates resolve unknown relationships and increase the proportion of pedigree known. For pedigrees that use analytical assumptions to base conservation decisions, genome-based relatedness estimates can confirm assumed relationships or prevent incorrect assumptions from plaguing conservation breeding programs, errors that would be exacerbated over multiple generations. More complete pedigrees with accurate relationships among individuals lead to higher overall pedigree confidence and pairing recommendations that minimize mean kinship. We acknowledge that mean kinship is one of many factors that inform conservation breeding pairing recommendations, including other genetic metrics (e.g., inbreeding), demography at institutions, health, and behavior of individuals (Lacy et al., 2012).

There was variable performance of relatedness estimators across species, sequencing method, degree of missing data, and pedigree knownness and depth. Thus, we advocate for explicit evaluation of multiple candidate relatedness estimators for each new dataset and system. Researchers can use our methods as a generalized workflow to test the relative accuracy and precision of relatedness estimators against a set of known pedigree relationships for their specific system. While this approach leverages known relationship classes to evaluate relatedness estimators, we appreciate this may not be feasible for all systems, especially wild systems without pedigree data. Some software programs are available to simulate data and compare estimators for relatedness with given datasets (e.g., COANCESTRY, or ‘related’; Wang et al., 2011; Pew et al., 2015). However, few can simulate large numbers of SNPs and no comparative software currently available includes relatedness estimators created for large SNP datasets. We therefore advocate for the investment in simulation software that can compare new genome-based relatedness estimators to evaluate estimator effectiveness. In the meantime, our results lead to a few general recommendations based on the sequencing method used, the amount of missing data in the system, and the availability of reliable pedigree data. The most accurate estimates of relatedness are predicted to come from genomic datasets generated using sequencing approaches that maximize statistical power (e.g., WGS) with nominal missing data (e.g., high-quality samples, single batch sequencing). We recommend using WGS data over RRS, when it is not cost, logistically, or computationally prohibitive, and further advocate for high quality DNA sampling, to provide the best possible sequencing reads. When pedigree information is unavailable for comparison, particular care must be used to ensure sufficient data quality. Lastly, even a robust relatedness estimator cannot overcome sampling deficiencies, so we urge proactive sample collection across the breadth and depth of pedigrees to facilitate accurate reconstruction of relationships among individuals to improve pedigree-based management. Adding newly resolved genome-based relatedness to species’ pedigrees uses more sources of evidence than using pedigree kinship alone and ultimately provides deeper insight into relationships within the conservation breeding population. If a new baseline of kinships utilizing genomic data can be established, breeding recommendations can be improved which will assist in the long-term preservation of genetic diversity.

## Acknowledgments

We are grateful for the continued support of Te Rūnanga o Ngāi Tahu, Te Ngāi Tūāhuriri Rūnanga, Te Rūnanga o Arowhenua, Te Rūnanga o Waihao and Te Rūnanga o Moeraki. We thank all members of the New Zealand Department of Conservation’s Kakī and Kākāriki Karaka Recovery Programmes, including the Isaac Conservation and Wildlife Trust, for their ongoing support. The Kakī and Kākāriki Karaka research was funded by the Ministry of Business, Innovation and Employment Endeavour Fund (UOCX1602 awarded to TES), the Brian Mason Scientific and Technical Trust (awarded to SJG and TES), and the Mohua Charitable Trust (awarded to TES). We would like to thank the Institute of Museum and Library Services National Leadership, Grant MG-30-15-0102-15 awarded to EKL. Support for SSH and SJG came from University of Wisconsin-Milwaukee College of Letters and Sciences, and a NSF EPSCoR RII Track-2 award (OIA-1826801), respectively. Special thanks to the AZA institutions that supplied samples: Buffalo Zoo, Brookfield Zoo, Dallas Zoo, Fossil Rim Wildlife Center, Frozen Zoo at the San Diego Zoo Wildlife Alliance, Kansas City Zoo, Louisville Zoo, Cleveland Metroparks Zoo, Omaha’s Henry Doorly Zoo and Aquarium, The Living Desert Zoo and Gardens, San Diego Zoo, and Saint Louis Zoo. Lastly, thanks to J. Ivy and Association of Zoos and Aquariums’ Molecular Data for Population Management Scientific Advisory Group who facilitated this collaboration.

## Data Accessibility

All R scripts necessary to replicate data analyses will be available on Dryad upon publication. For Addax, Koala, and Inca Tern, species managed by the Association of Zoos and Aquariums, genomic data will be available in VCF format upon acceptance on Dryad.

A subset of the genomic data used in this project were previously derived from two culturally significant species that are taonga (treasured) to Māori (the Indigenous Peoples of Aotearoa New Zealand): Kakī/Black Stilt and Kākāriki Karaka/Orange-Fronted Parakeet (Galla et al., 2019; Galla et al., 2020). For Māori, all genomic data obtained from taonga species have whakapapa (genealogy that includes people, plants and animals, mountains, rivers and winds) and are therefore taonga in their own right (Collier-Robinson et al., 2019; Hudson et al. 2022). Thus, these data are tapu (sacred), and tikanga (customary practices, protocols, and ethics) determine how people interact with them (Collier-Robinson et al., 2019; Hudson et al. 2022). To this end, genomics data for Kakī and Kākāriki Karaka will be made available from a local genome browser (http://www.ucconsert.org/data/) on the recommendation of the kaitiaki (guardians) for the iwi (tribes) and hapū (subtribes) that affiliate with them.

## Benefit Sharing Statement

We present genomic data for Kakī and Kākāriki Karaka consistent with both FAIR and CARE data principles (Carroll et al., 2020, 2021; McCartney et al., 2022). Our research findings have been used to inform the conservation breeding programs herein and are broadly applicable to management of threatened species around the globe. This publication is the result of international collaboration between scientists at research institutions in the United States and New Zealand, all of whom are named co-authors on this publication.

## Authors’ Contributions

SSH, SJG, EKL, and TES conceived the research ideas and designed the methodology. SSH and SJG generated the genomic datasets. SSH analyzed the data. SSH and SJG led the writing of the manuscript. EKL, TES, and ASP supervised the analysis and interpretation of the research. All authors contributed to the manuscript preparation and gave final approval for submission.

## Supplemental Figure Captions

**Figure S1.**
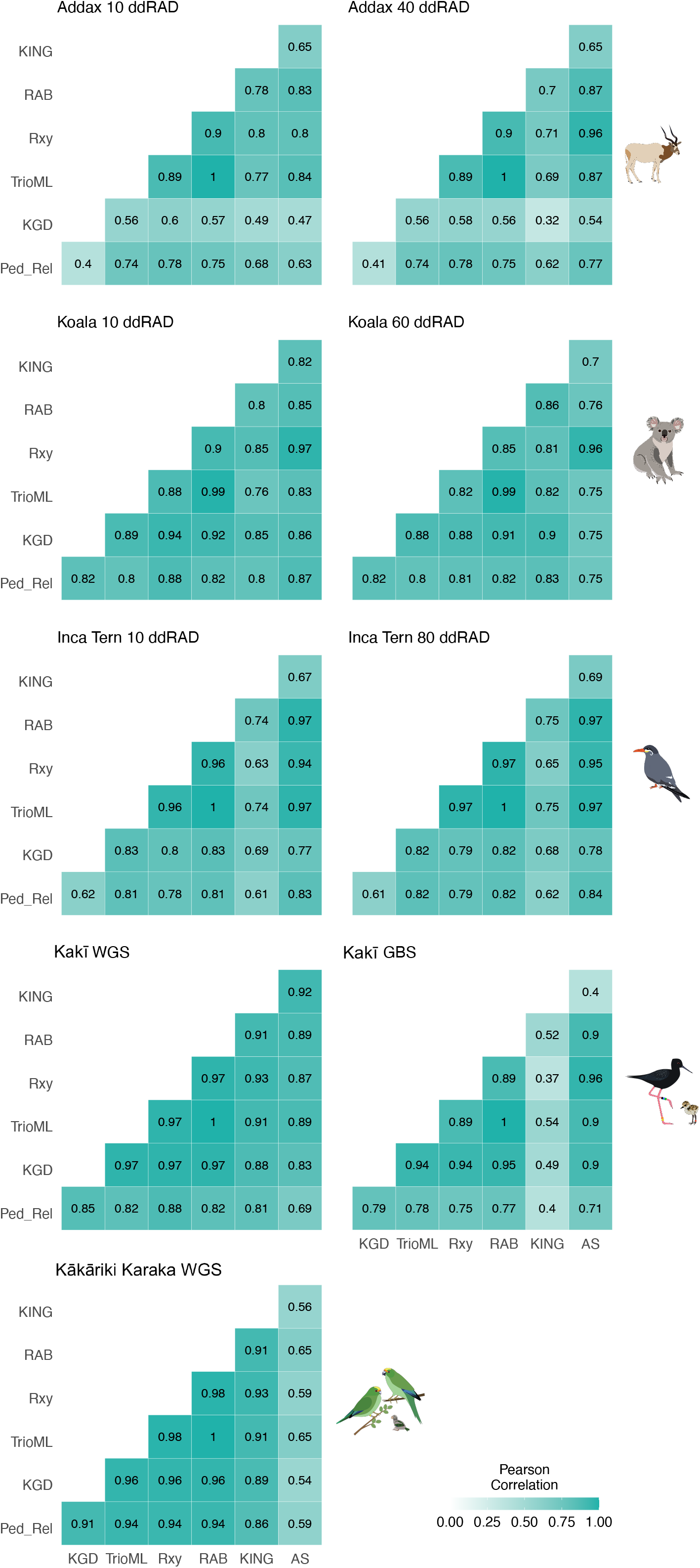
Heatmaps depicting pairwise Pearson’s correlation coefficients between all six relatedness estimators (kinship genetic distance (KGD), Wang Maximum Likelihood (TrioML), Queller and Goodnight (R_xy_), Kinship INference for Genome-wide association studies (KING-robust), and Pairwise Relatedness (RAB), allele-sharing co-ancestry (AS)) for each of the 9 genomic datasets: Addax ddRAD data with max 10% missing data (Addax 10), Addax ddRAD data with max 40% missing data (Addax 40), Koala ddRAD data with 10% max missing data (Koala 10), Koala ddRAD data with 60% max missing data (Koala 60), Inca Tern ddRAD data with 10% max missing data (Inca Tern 10), Inca Tern ddRAD data with 80% max missing data (Inca Tern 80), Kakī WGS data, Kakī GBS data, and Kākāriki Karaka WGS data. Darker gradient colors denote higher correlation values.

## Estimators of Relatedness

Below are descriptions of kinship and relatedness estimators compared in this study.

### Kinship Genetic Distance (KGD)

KGD is a method of moments estimator of the genomic relatedness matrix (GRM) that was designed for genotyping-by-sequencing data but can be used broadly with RADseq or resequencing data. Allele frequencies of the sampled population are used as a reference to estimate the correlation between pairwise genotypes across all loci. KGD improves this relatedness estimation by incorporating an adjustment based on sequencing depth (Dodds et al 2015). While estimates of self-relatedness are improved when sequencing depth is between 5 - 10, SNPs sequenced at lower depth (~1 - 4) between pairs of individuals may be used with confidence. This method capitalizes on the tradeoff between the number of SNPs analyzed and the depth of the SNPs retained to optimize the dataset for analyses.

### Wang Maximum Likelihood (TrioML)

TrioML is a triadic likelihood method that calculates the probability of individuals falling into a particular relationship category. The estimates of relatedness between pairs of individuals are in relation to a third reference individual to reduce errors arising from allelic identity by state rather than identity by descent (Wang, 2007, 2011). Reference allele frequencies from populations in Hardy-Weinberg equilibrium are required for TrioML estimation. The estimator is bounded by 0 and 1, and is considered robust to inbreeding, small sampling sizes, and genotyping error.

### Queller and Goodnight (R_xy_)

R_XY_ is a method of moments approach to relatedness that is estimated based on allele frequency differences between pairs of individuals from the population mean (Queller and Goodnight, 1989). Reference allele frequencies are calculated after excluding data from the pair of individuals being analyzed. The estimator is considered statistically unbiased, thus ranges from −1 to 1, and requires scaling or truncating the negative numbers to be applied as biologically relevant relatedness estimates that range from 0 to 1. Both R_xy_ and TrioML are considered to have the best performance in estimating 1^st^ order relationships compared to more distant relationships (Csillery et al. 2006).

### Pairwise Relatedness (R_AB_)

R_AB_ is a relatedness estimator first described in Hedrick and Lacy 2014 (where it was referred to as r_xy_) that is calculated from the shared proportion of homologous alleles identity by descent using the subset of the nine possible genotypes between two individuals at a biallelic locus. The estimator has an upper bound of 1 so that self-relatedness is always 1 irrespective of inbreeding. Because of this, R_AB_ is expected to be less than the coefficient of relatedness (2*kinship) when inbreeding is present in one or both individuals. Similar to KING-robust, when implemented in NgsRelate V2 (Hanhgoj et al. 2019), genotype likelihoods are used to improve accuracy in RADseq or resequencing data. It should be noted that NgsRelate V2 uses reference allele frequencies to calculate genotype likelihoods for R_AB_.

### Kinship INference for Genome-wide association studies (KING-robust)

Pairwise kinships are estimated without reference allele frequencies, but through estimating the identity by state (IBS) probabilities that are compared to expectations for different relationship categories (e.g., full sibling, half sibling; Manichaikul et al., 2010; Waples et al. 2019) in the presence of population structure. The estimator is based on a subset of the nine possible genotype combinations between two individuals at a biallelic locus in non-inbred individuals. KING is commonly implemented with the software program ANGSD and a subprogram within it, NgsRelate V2 which uses genotype likelihoods and not called genotypes to reduce bias when sequencing depth is low in RADseq or resequencing data (Hangoj et al. 2019). While allele frequencies are not used in the NgsRelate V2 estimate of KING-robust kinship, the program does use a two-dimensional site frequency spectrum (2D-SFS) derived from the genotype likelihoods.

### Allele-Sharing (AS)

Allele-sharing is a similarity index, often referred to as molecular coancestry. It is the probability that an allele chosen randomly from a given locus in two different individuals is identical by state, irrespective of whether the alleles are inherited from a recent common ancestor (Caballero and Toro 2002, Eding and Meuwissen 2001). Because pairwise allele sharing values are typically high (>0.6), especially when populations are inbred, or large numbers of biallelic loci with low minor allele frequencies are used, they are not directly comparable to pedigree-based kinships and require scaling to be used as such. This estimation differs from the relatedness coefficient that estimates the proportion of shared alleles so that a pair of individuals share 0, 1, or 2 alleles by descent at each genomic locus. See KING-robust, R_AB_.

## References

Attard, C. R., Beheregaray, L. B., & Möller, L. M. (2018). Genotyping □by□ sequencing for estimating relatedness in nonmodel organisms: Avoiding the trap of precise bias. Molecular ecology resources, 18(3), 381–390.

Ballou, J. D., & Lacy, R. C. (1995). Identifying genetically important individuals for management of genetic variation in pedigreed populations. Population management for survival and recovery, 76–111.

Balloux, F., Amos, W., & Coulson, T. (2004). Does heterozygosity estimate inbreeding in real populations? Molecular Ecology, 13(10), 3021–3031.

Baruch, E., & Weller, J. I. (2008). Estimation of the number of SNP genetic markers required for parentage verification. Animal Genetics, 39(5), 474–479.

Bergner, L. M., Jamieson, I. G., & Robertson, B. C. (2014). Combining genetic data to identify relatedness among founders in a genetically depauperate parrot, the Kakapo (Strigops habroptilus). Conservation Genetics, 15(5), 1013–1020.

Carroll, S. R., Garba, I., Figueroa-Rodríguez, O. L., Holbrook, J., Lovett, R., Materechera, S., Parsons, M., raseroka, K., Rodriguez-Lonebear, D., Rowe, R., Sara, R. Walker, J.D., Anderson, J., & Hudson, M. (2020). The CARE Principles for Indigenous Data Governance.

Carroll, S. R., Herczog, E., Hudson, M., Russell, K., & Stall, S. (2021). Operationalizing the CARE and FAIR Principles for Indigenous data futures. Scientific Data, 8(1), 1–6.

Cassell, B. G., Adamec, V., & Pearson, R. E. (2003). Effect of incomplete pedigrees on estimates of inbreeding and inbreeding depression for days to first service and summit milk yield in Holsteins and Jerseys. Journal of Dairy Science, 86(9), 2967–2976.

Catchen, J., Hohenlohe, P. A., Bassham, S., Amores, A., & Cresko, W. A. (2013). Stacks: An analysis tool set for population genomics. Molecular Ecology, 22(11), 3124–3140.

Csilléry, K., Johnson, T., Beraldi, D., Clutton-Brock, T., Coltman, D., Hansson, B.,… & Pemberton, J. M. (2006). Performance of marker-based relatedness estimators in natural populations of outbred vertebrates. Genetics, 173(4), 2091–2101.

Chemnick, L. G., Houck, M. L., & Ryder, O. A. (2009). Banking of Genetic Resources. Conservation genetics in the age of genomics (pp. 124–130).

Clarke, A. G. (2009). The Frozen Ark Project: The role of zoos and aquariums in preserving the genetic material of threatened animals. International Zoo Yearbook, 43(1), 222–230. Retrieved June 9, 2021, from http://www.frozenark.org

Comizzoli, P. (2014). C-29: The Pan-Smithsonian cryo-initiative-freezing for the future. Cryobiology, 69(3), 509.

Danecek, P., Auton, A., Abecasis, G., Albers, C. A., Banks, E., DePristo, M. A., Handsaker, R. E., et al. (2011). The variant call format and VCFtools. Bioinformatics, 27(15), 2156–2158.

Dodds, K. G., McEwan, J. C., Brauning, R., Anderson, R. M., Stijn, T. C., Kristjánsson, T., & Clarke, S. M. (2015). Construction of relatedness matrices using genotyping-by-sequencing data. BMC Genomics, 16(1), 1047.

Enright W. (2019) International Addax Studbook. Gland Switzerland: World. Association of Zoos & Aquariums.

Faust, L., Bergstrom, Y., Thompson, S., & Bier, L. (2019). PopLink Version 2.5. Lincoln Park Zoo. Chicago, IL.

Fernández, J., Villanueva, B., Pong-Wong, R., & Toro, M. Á. (2005). Efficiency of the use of pedigree and molecular marker information in conservation programs. Genetics, 170(3), 1313–1321.

Ferrie, G. (2017). Using Molecular Genetic and Demographic Tools to Improve Management of Ex Situ Avian Populations.

Flanagan, A. M., Masuda, B., Grueber, C. E., & Sutton, J. T. (2021). Moving from trends to benchmarks by using regression tree analysis to find inbreeding thresholds in a critically endangered bird. Conservation Biology.

Flanagan, S. P., & Jones, A. G. (2019). The future of parentage analysis: From microsatellites to SNPs and beyond. Molecular Ecology, 28(3), 544–567.

Galla, S. J. (2019) Conservation genomic management of two critically-endangered New Zealand birds. Doctoral Thesis, University of Canterbury. http://dx.doi.org/10.26021/8081

Galla, S. J., Forsdick, N. J., Brown, L., Hoeppner, M. P., Knapp, M., Maloney, R. F., Moraga, R., et al. (2019). Reference genomes from distantly related species can be used for discovery of single nucleotide polymorphisms to inform conservation management. Genes, 10(1).

Galla, S. J., Moraga, R., Brown, L., Cleland, S., Hoeppner, M. P., Maloney, R. F., Richardson, A., et al. (2020). A comparison of pedigree, genetic and genomic estimates of relatedness for informing pairing decisions in two critically endangered birds: Implications for conservation breeding programmes worldwide. Evolutionary Applications, 13(5), 991–1008.

Galla, S. J., Brown, L., Couch-Lewis, Y. Steeves, T. E. (2021). The relevance of pedigrees in the conservation genomics era. Molecular Ecology, doi: https://doi.org/10.1111/mec.16192,

Giglio, R. M., Ivy, J. A., Jones, L. C., & Latch, E. K. (2016). Evaluation of alternative management strategies for maintenance of genetic variation in wildlife populations. Animal Conservation, 19(4), 380–390.

Giglio, Rachael M., Ivy, J. A., Jones, L. C., & Latch, E. K. (2018). Pedigree-based genetic management improves bison conservation. Journal of Wildlife Management, 82(4), 766–774.

Glaubitz, J. C., Casstevens, T. M., Lu, F., Harriman, J., Elshire, R. J., Sun, Q., & Buckler, E. S. (2014). TASSEL-GBS: A High Capacity Genotyping by Sequencing Analysis Pipeline. PLoS ONE, 9(2), e90346.

Goodnight, K. F., Queller, D. C., & Goodnight, K. F. (1999). Computer software for performing likelihood tests of pedigree relationship using genetic markers. Molecular Ecology, 8(7), 1231–1234.

Gooley, R., Hogg, C. J., Belov, K., & Grueber, C. E. (2017). No evidence of inbreeding depression in a Tasmanian devil insurance population despite significant variation in inbreeding. Scientific Reports, 7(1).

Goudet, J., Kay, T., & Weir, B. S. (2018). How to estimate kinship. Molecular ecology, 27(20), 4121–4135.

Graham, C. F., Glenn, T. C., Mcarthur, A. G., Boreham, D. R., Kieran, T., Lance, S., Manzon, R. G., et al. (2015). Impacts of degraded DNA on restriction enzyme associated DNA sequencing (RADSeq). Molecular Ecology Resources, 15(6), 1304–1315.

Gutiérrez, J. P., Royo, L. J., Álvarez, I., & Goyache, F. (2005). MolKin v2.0: A computer program for genetic analysis of populations using molecular coancestry information. Journal of Heredity, 96(6), 718–721.

Hamlin C. (2019) AZA Regional Queensland Koala (Phascolarctos cinereus) Studbook. Maryland: Association of Zoos & Aquariums.

Hammerly, S. C., de la Cerda, D. A., Bailey, H., & Johnson, J. A. (2016). A pedigree gone bad: increased offspring survival after using DNA-based relatedness to minimize inbreeding in a captive population. Animal Conservation, 19(3), 296–303.

Hanghøj, K., Moltke, I., Andersen, P. A., Manica, A., & Korneliussen, T. S. (2019). Fast and accurate relatedness estimation from high-throughput sequencing data in the presence of inbreeding. Gigascience, 8(5), giz034.

Hedrick, P. P. W., Lacy, R. C., & Baker, C. S. (2015). Measuring relatedness between inbred individuals. Journal of Heredity, 106(1), 20–25.

Henkel, J. R., Jones, K. L., Hereford, S. G., Savoie, M. L., Leibo, S. P., & Howard, J. J. (2012). Integrating microsatellite and pedigree analyses to facilitate the captive management of the endangered Mississippi sandhill crane (Grus canadensis pulla). Zoo Biology, 31(3), 322–335.

Hogg, C. J., Wright, B., Morris, K. M., Lee, A. V., Ivy, J. A., Grueber, C. E., & Belov, K. (2019). Founder relationships and conservation management: empirical kinships reveal the effect on breeding programmes when founders are assumed to be unrelated. Animal Conservation, 22(4), 348–361.

Hudson, M., Battershill, C., Thompson, A., Scott, M., Wilcox, P., Brooks, R.T., Mika, J., and Warbrick, L. (2022). Te Nohonga Kaitiaki: Guidelines for Genomic Research with Maori. https://www.genomics-aotearoa.org.nz/education-resources/guidelines

Ivy, J. A., & Putnam, A. S. (2019). Calculate Allele Sharing Coefficients (CASC) (version 1.0). San Diego, CA, USA: San Diego Zoo Global.

Ivy, J. A., Putnam, A. S., Navarro, A. Y., Gurr, J., & Ryder, O. A. (2016). Applying SNP-derived molecular coancestry estimates to captive breeding programs. Journal of Heredity, 107(5), 403–412.

Jannink, J. L., Bink, M. C. A. M., & Jansen, R. C. (2001). Using complex plant pedigrees to map valuable genes. Trends in Plant Science, 6(8), 337–342.

Jones, O. R., & Wang, J. (2010). Molecular marker-based pedigrees for animal conservation biologists. Animal Conservation, 13(1), 26–34.

Korneliussen, T. S., & Moltke, I. (2015). NgsRelate: A software tool for estimating pairwise relatedness from next-generation sequencing data. Bioinformatics, 31(24), 4009–4011.

Lacy, R C, & Pollak, J. P. (2014). Vortex: A Stochastic Simulation of the Extinction Process. Manual. Version 9.99c.

Lacy, R.C., Ballou, J. D., & Pollak, J. P. (2012). PMx: Software package for demographic and genetic analysis and management of pedigreed populations. Methods in Ecology and Evolution, 3(2), 433–437.

Leigh, D. M., Lischer, H. E. L., Grossen, C., & Keller, L. F. (2018). Batch effects in a multiyear sequencing study: False biological trends due to changes in read lengths. Molecular Ecology Resources, 18(4), 778–788.

Lemopoulos, A., Prokkola, J. M., Uusi-Heikkilä, S., Vasemägi, A., Huusko, A., Hyvärinen, P., Koljonen, M. L., et al. (2019). Comparing RADseq and microsatellites for estimating genetic diversity and relatedness — Implications for brown trout conservation. Ecology and Evolution, 9(4), 2106–2120.

Li, H., Handsaker, B., Wysoker, A., Fennell, T., Ruan, J., Homer, N., Marth, G., et al. (2009). The Sequence Alignment/Map format and SAMtools. Bioinformatics, 25(16), 2078–2079.

McCartney, A. M., Anderson, J., Liggins, L., Hudson, M. L., Anderson, M. Z., TeAika, B., Geary, J., Cook-Deegan, R., Patel, H.R., & Phillippy, A. M. (2022). Balancing openness with Indigenous data sovereignty: An opportunity to leave no one behind in the journey to sequence all of life. Proceedings of the National Academy of Sciences, 119(4).

Milligan, J. L., Davis, A. K., & Altizer, S. M. (2003). Errors associated with using colored leg bands to identify wild birds. Journal of Field Ornithology, 74(2), 111–118.

Montgomery, M. E., Ballou, J. D., Nurthen, R. K., England, P. R., Briscoe, D. A., & Frankham, R. (1997). Minimizing kinship in captive breeding programs. Zoo Biology, 16(5), 377–389.

Morin, P. A., Martien, K. K., & Taylor, B. L. (2009). Assessing statistical power of SNPs for population structure and conservation studies. Molecular Ecology Resources, 9(1), 66–73.

Mucha, S., & Windig, J. J. (2009). Effects of incomplete pedigree on genetic management of the Dutch Landrace goat. Journal of Animal Breeding and Genetics, 126(3), 250–256.

Nelson S. (2017) AZA Regional Inca Tern (Larosterna inca) Studbook. Maryland: Association of Zoos & Aquariums.

Nelson S., Lynch C. (2021) Population Analysis & Breeding and Transfer Plan, Inca Tern (Larosterna inca) Species Survival Plan. Maryland: Association of Zoos & Aquariums

Ott, J. (1974). Estimation of the recombination fraction in human pedigrees: efficient computation of the likelihood for human linkage studies. American Journal of Human Genetics, 26(5), 588–597.

Overbeek, A., Galla, S. J., Brown, L., Cleland, S., Thyne, C., Maloney, R., & Steeves, T. E. (2020). Pedigree validation using genetic markers in an intensively managed taonga species, the critically endangered kakī (Himantopus novaezelandiae). Notornis, 67, 709–716.

Paris, J. R., Stevens, J. R., & Catchen, J. M. (2017). Lost in parameter space: a road map for stacks. Methods in Ecology and Evolution, 8(10), 1360–1373.

Pemberton, J. M. (2008). Wild pedigrees: the way forward. Proceedings of the Royal Society B: Biological Sciences, 275, 613–621.

Peterson, B. K., Weber, J. N., Kay, E. H., Fisher, H. S., & Hoekstra, H. E. (2012). protocol to ddRAD. PLoS ONE.

Pew, J., Muir, P. H., Wang, J., & Frasier, T. R. (2015). related: an R package for analysing pairwise relatedness from codominant molecular markers. Molecular ecology resources, 15(3), 557–561.

Putnam, A. S., & Ivy, J. A. (2014). Kinship-based management strategies for captive breeding programs when pedigrees are unknown or uncertain. Journal of Heredity, 105(3), 303–311.

Queller, D. C., & Goodnight, K. F. (1989). Estimating relatedness using genetic markers. Evolution, 43(2), 258–275.

R Core Team (2018). R: A language and environment for statistical computing. R Foundation for Statistical Computing, Vienna, Austria. URL https://www.R-project.org/.

Robinson, S. P., Simmons, L. W., & Kennington, W. J. (2013). Estimating relatedness and inbreeding using molecular markers and pedigrees: the effect of demographic history. Molecular Ecology, 22, 5779–5792.

Rudnick, J. A., & Lacy, R. C. (2008). The impact of assumptions about founder relationships on the effectiveness of captive breeding strategies. Conservation Genetics, 9(6), 1439–1450.

Sanders, M. D., & Maloney, R. F. (2002). Causes of mortality at nests of ground-nesting birds in the Upper Waitaki Basin, South Island, New Zealand: a 5-year video study. Biological conservation, 106(2), 225–236.

Santure, A. W., Stapley, J., Ball, A. D., Birkhead, T. R., Burke, T., & Slate, J. (2010). On the use of large marker panels to estimate inbreeding and relatedness: Empirical and simulation studies of a pedigreed zebra finch population typed at 771 SNPs. Molecular Ecology, 19(7), 1439–1451.

Skare, Ø., Sheehan, N., & Egeland, T. (2009). Identification of distant family relationships. Bioinformatics, 25(18), 2376–2382.

Smouse, P. E. (2010). How many SNPs are enough? Molecular Ecology.

Sonesson, A. K., & Meuwissen, T. H. E. (2001). Minimization of rate of inbreeding for small populations with overlapping generations. Genetical Research, 77(3), 285–292.

Spielman, D., Brook, B. W., Briscoe, D. A., & Frankham, R. (2004). Does inbreeding and loss of genetic diversity decrease disease resistance? Conservation Genetics, 5(4), 439–448.

Spielman, D., & Frankham, R. (1992). Modeling problems in conservation genetics using captive Drosophila populations: Improvement of reproductive fitness due to immigration of one individual into small partially inbred populations. Zoo Biology, 11(5), 343–351.

Templeton, A. R., & Read, B. (1994). Inbreeding: one word, several meanings, much confusion. In Conservation genetics (pp. 91–105).

Sun, M., Jobling, M. A., Taliun, D., Pramstaller, P. P., Egeland, T., & Sheehan, N. A. (2016). On the use of dense SNP marker data for the identification of distant relative pairs. Theoretical Population Biology, 107, 14–25.

Wang, J. (2007). Parentage and sibship exclusions: Higher statistical power with more family members. Heredity, 99(2), 205–217.

Wang, J. (2014). Marker-based estimates of relatedness and inbreeding coefficients: An assessment of current methods. Journal of Evolutionary Biology, 27(3), 518–530.

Wang, Jinliang. (2002). An estimator for pairwise relatedness using molecular markers. Genetics, 160(3), 1203–1215.

Wang, Jinliang. (2011). Coancestry: A program for simulating, estimating and analysing relatedness and inbreeding coefficients. Molecular Ecology Resources, 11(1), 141–145.

Waples, R. K., Albrechtsen, A., & Moltke, I. (2019). Allele frequency-free inference of close familial relationships from genotypes or low-depth sequencing data. Molecular Ecology, 28(1), 35–48.

Weeks, A. R., Sgro, C. M., Young, A. G., Frankham, R., Mitchell, N. J., Miller, K. A., Byrne, M., et al. (2011). Assessing the benefits and risks of translocations in changing environments: A genetic perspective. Evolutionary Applications, 4(6), 709–725.

Wickham, H. (2011). ggplot2. Wiley Interdisciplinary Reviews: Computational Statistics, 3(2), 180–185.

Wildt, D., Miller, P., Koepfli, K. P., Pukazhenthi, B., Palfrey, K., Livingston, G., Beetem, D., et al. (2019). Breeding Centers, Private Ranches, and Genomics for Creating Sustainable Wildlife Populations. BioScience.

Wolfe, B. A., Aguilar, R. F., Aguirre, A. A., Olsen, G. H., & Blumer, E. S. (2012). Sorta situ: The new reality of management conditions for wildlife populations in the absence of “Wild” spaces. New Directions in Conservation Medicine: Applied Cases of Ecological Health.

Wright, S. (1922). Coefficients of Inbreeding and Relationship. The American Naturalist, 56(645), 330–338.

